# Single-molecule microscopy reveals that TFIIE subunits dynamically interact with preinitiation complexes in a manner controlled by TFIIH

**DOI:** 10.1101/2025.04.19.649533

**Authors:** Stephen R. Archuleta, Ryan C. Miller, Julia A. Mirita, James A. Goodrich, Jennifer F. Kugel

## Abstract

Transcription by RNA polymerase II (Pol II) requires general transcription factors that bind with Pol II at the promoters of protein-coding genes to form preinitiation complexes (PICs). Among these is TFIIE, which recruits TFIIH to the PIC and stimulates the kinase and translocase activities of TFIIH, thereby regulating the fate of formed PICs. In this study, we used a purified reconstituted human Pol II transcription system and single molecule total internal reflection fluorescence (smTIRF) microscopy to monitor TFIIE binding dynamics in PICs under different conditions in real time. We observed highly dynamic interactions of the two subunits of TFIIE (TFIIEα and TFIIEβ) with PICs. Measurement of rate constants for on/off binding of each subunit suggest they behave asynchronously. TFIIH exclusion increased the rates of association and dissociation for both subunits, with the strongest effect on TFIIEα. Despite stabilization of TFIIE by TFIIH the TFIIE subunits remain dynamic in PICs. Additionally, two disease-related TFIIEβ point mutations destabilized TFIIEβ and altered its kinetic behaviors within PICs. Our results contribute to an emerging model that PICs are not static assemblies and highlight important connections between the structural arrangement and kinetic behaviors of GTFs in PICs.

## Introduction

Transcription by RNA polymerase II (Pol II) is a highly complex process that requires the coordination of an array of general and regulatory factors that act throughout the transcription reaction. The general transcription factors (GTFs) are machinery used to transcribe all genes by Pol II, with critical roles during the early stages of transcription: preinitiation complex (PIC) assembly, promoter melting, initiation, and promoter escape. The GTFs (TFIID, TFIIA, TFIIB, TFIIF, TFIIE, and TFIIH) assemble with Pol II on promoter DNA to form PICs [1,2]. Extensive biochemical research has described general mechanisms for PIC assembly, including stepwise and holoenzyme models [2,3]. In the stepwise mechanism of assembly, the GTFs assemble in the order listed above, with Pol II binding to the PIC with TFIIF. Each factor is believed to facilitate the recruitment of the next factor in the sequence [3–5]. In the holoenzyme model of assembly, Pol II and many of the GTFs pre-assemble away from the DNA, then bind as a subcomplex to the core promoter [2,3,6]. These mechanisms are not mutually exclusive, and it is likely both occur in cells as unique ways to regulate transcriptional activity in response to different cellular stimuli.

In recent years, the architecture of PICs has been revealed via intricate cryoEM structures [7–12]. CryoEM allows multiple conformations of a complex to be resolved to accommodate flexibility; however, the structural pictures of transcription complexes obtained from cryoEM are largely static. Real-time single molecule fluorescence imaging has illuminated dynamic mechanisms of Pol II assembly and transcription, both in cells and in vitro. In cells, Pol II has been visualized via super-resolution fluorescence microscopy, revealing phenomena such as transcriptional bursting, Pol II clustering, and formation of condensates [13–20]. Single-particle tracking of GTFs and Pol II in live yeast cells revealed that PIC assembly and initiation-coupled disassembly is a dynamic process that occurs over seconds [21]. In vitro, single-molecule studies of Pol II and GTFs have revealed unexpectedly dynamic modes of PIC assembly and disassembly during RNA synthesis [22–24]. The kinetics governing assembly of GTFs in PICs, and their dynamic behaviors during disassembly, are not yet fully understood. Learning how GTFs associate and dissociate from the PIC and how the GTFs impact one another is important for uncovering mechanisms of transcriptional regulation.

TFIIE, one of the Pol II GTFs, consists of a heterodimer of TFIIEα and TFIIEβ subunits, with masses of 56 kDa and 34 kDa, respectively. TFIIE serves two primary purposes during early transcription: 1) recruitment of TFIIH to the PIC, which completes PIC assembly in the stepwise model, and 2) stimulation of the ATP-dependent kinase and translocase activities of TFIIH [25–28]. The kinase activity of the CDK7 subunit of TFIIH phosphorylates the long, unstructured C-terminal tail (CTD) of the Rpb1 subunit of Pol II on the Ser5 and Ser7 residues in the heptapeptide repeat (consensus YSPTSPS) [29–31]. These phosphorylation events help transition Pol II into different phases of the transcription cycle and create a binding platform for co-transcriptional factors [29,30]. The translocase activity of the XPB subunit of TFIIH uses contacts downstream of the transcription start site (around +20) to facilitate promoter melting using mechanical and torsional strain [10,32–37]. TFIIEα stabilizes DNA in the PIC before promoter melting [34,38], while TFIIEβ stabilizes closed and open DNA conformations through a network of winged-helix motifs, in conjunction with TFIIF [34,35,39–41]. After initiation takes place, TFIIE has also been shown to facilitate promoter clearance and the transition from initiation to elongation [42–44].

Data in recent years has suggested that TFIIE exhibits unique behavior in PICs beyond the recruitment of TFIIH. For example, a study using a reconstituted human transcription system and quantitative analysis of Western blots observed that the TFIIE subunits release from PICs independently, with TFIIEα releasing after phosphorylation of the Pol II CTD and TFIIEβ after initiation [45]. A single-molecule study investigating activator-dependent PIC assembly in yeast extracts found that Tfa2 (yeast analog for TFIIEβ) remained highly dynamic in PICs even after the incorporation of TFIIH [23]. This dynamic behavior was also seen in single molecule tracking experiments in yeast cells with labeled Tfa1 (TFIIEα analog) [21]. Additionally, cryoEM studies have noted difficulty resolving TFIIE within the PIC when TFIIH was excluded, suggesting that TFIIH binding stabilizes the binding of TFIIE [35,38]. These studies support a dynamic role for TFIIE within PICs, but much remains to be learned about how the TFIIE subunits behave and are regulated.

Here we used single molecule total internal reflection fluorescence (smTIRF) microscopy to investigate the dynamic interactions of TFIIE subunits in PICs in real time. Our experiments used a highly purified reconstituted human Pol II transcription system with fluorescently labeled TFIIEα and TFIIEβ to measure rate constants governing TFIIE association and dissociation from PICs. We observed rapid on/off binding behaviors for both TFIIE subunits that suggest asynchronous behavior. TFIIH exclusion increased the frequency of on/off binding for both subunits, and their rates of association and dissociation, with TFIIEα showing a much stronger destabilization. This suggests that TFIIH stabilizes TFIIE in PICs, although TFIIE subunits remain dynamic even with TFIIH present. Additionally, we determined that two disease-relevant TFIIEβ point mutants (A150P and D187Y) altered the kinetic behaviors of TFIIEβ. Together, our findings contribute to the emerging view that PICs are not static structures and give insight into how the structural arrangement of the PIC influences the dynamic interactions of TFIIE.

## Results

### A single molecule system for studying TFIIE subunits interacting with preinitiation complexes in real time

To visualize TFIIE subunits in our single molecule TIRF system, each was labeled with Alexa Fluor 647 (AF647) red fluorescent dye. TFIIEα contained a C-terminal SNAP-tag for labeling (TFIIEα-647). TFIIEβ naturally contains a unique surface-exposed cysteine residue that we labeled using maleimide chemistry (TFIIEβ-647). The labeled TFIIE subunits were tested using in vitro transcription assays with highly purified human proteins (TBP subunit of TFIID, TFIIB, TFIIF, TFIIE, TFIIH, and Pol II). The DNA template contained a core promoter upstream of a G-less cassette that allowed an RNA of defined length to be transcribed in the presence of ATP, CTP, and UTP [46,47]. As shown in Figure 1A, TFIIEα-647 and TFIIEβ-647 (red font) had activity similar to the unlabeled proteins, whether tested with their unlabeled counterpart subunit (lanes 6 and 7) or together (lane 8). The transcription system was dependent on both TFIIE subunits and TFIIH for activity (lanes 2-5).

**Figure 1.**
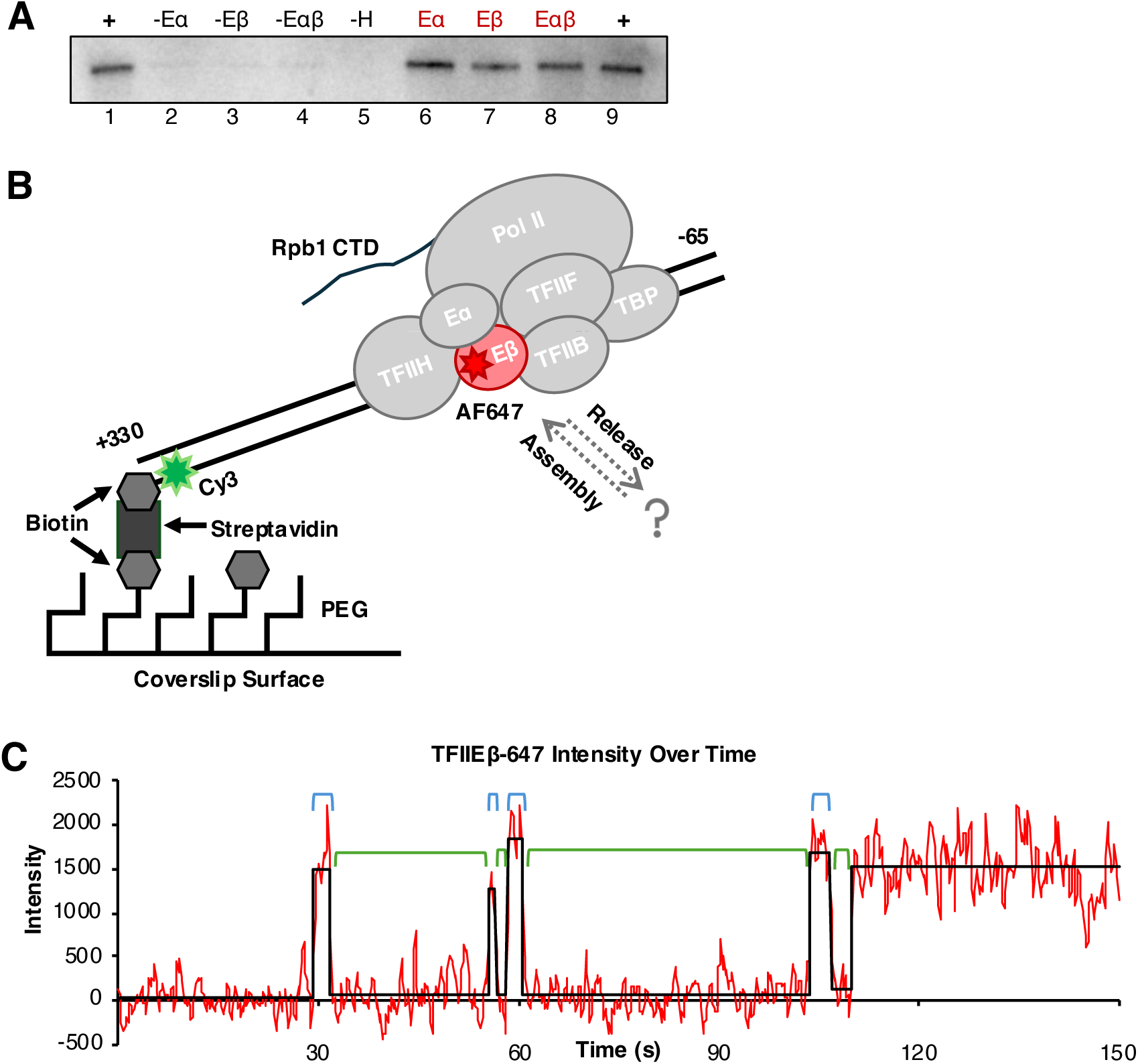
A single molecule system to measure the on/off binding of TFIIE subunits to PICs. A) AF647-labeled TFIIE subunits support transcription in vitro and are necessary for activity. Shown are ^32^P-labeled transcripts. B) The Cy3-labeled DNA template was immobilized on the slide surface using biotin-streptavidin interactions. Pol II and the GTFs assembled on the core promoter to form the PIC. One TFIIE subunit was labeled with AF647 (red). After PIC assembly, green and red images were collected to respectively capture DNA location and TFIIE dynamics over time. C) Representative trace showing a single molecule of TFIIEβ-647 binding and releasing from a single DNA over a time course of 150 seconds. Red line is emission from AF647, black line shows state changes detected by our software, blue brackets represent bound dwell times, and green brackets represent unbound dwell times. For display, data were smoothed using the moving boxcar average with a window size of 3.

After verifying their transcriptional activity, the fluorescently labeled TFIIE subunits were used in smTIRF experiments to test whether their association with PICs was dynamic or stable. The system used is diagrammed in Figure 1B. A coverslip was functionalized with a mixture of mPEG and biotin-PEG, and streptavidin-biotin interactions were used to immobilize the Cy3-labeled (green star) DNA template on the slide surface. To investigate the dynamics of each TFIIE subunit in PICs, experiments were performed as follows:

1) minimal PICs containing DNA, TBP, TFIIB, TFIIF, and Pol II were assembled and immobilized on the surface, 2) TFIIEα, TFIIEβ and TFIIH (with one TFIIE subunit labeled with AF647 (red star)) were flowed into the slide chamber, and 3) emissions in the green and red channels were imaged over time to capture DNA and TFIIE locations, respectively. PICs were identified on the slide surface by colocalizing green and red spots across the two movies. For each spot pair, we analyzed the behavior of the labeled TFIIE subunit over time, tracking changes in states (bound versus unbound). An example intensity trace of TFIIEβ-647 (red) is shown in Figure 1C; multiple state changes (black) were observed, reflecting dynamic behavior. The length of time a TFIIE subunit remained bound was categorized as a bound dwell time (blue brackets) and the time between two binding events was categorized as an unbound dwell time (green brackets). Dwell times were plotted and fit with exponential equations to obtain kinetic parameters, as described in subsequent figures. In control experiments, we assessed photobleaching using immobilized DNA labeled with AF647. After 2.5 minutes of imaging, the length of all of our experiments with TFIIE-647, 99% of spots remained; therefore, we are confident that loss of red signal due to photobleaching (as opposed to protein release) occurred rarely in our experiments.

For each experiment we performed a control to evaluate the colocalization of green and red movies that identified PICs. We rotated the Cy3 DNA movie 90° counterclockwise and colocalized it with the TFIIE-647 movie, as diagrammed in Figure 2A; spot pairs found with the rotated control do not represent PICs. We quantified dwell times from any spot pairs found in the rotated control. When compared to the normal PIC pairing, the rotated control pairing showed a sharp decrease in the number of dwell times (Figure 2B). In addition, we performed control experiments that excluded Pol II from PIC assembly. In these data sets, the number of dwell times found were the same level as the rotated control (Figure 2B), showing that the dynamic association of the TFIIE subunits with PICs is dependent on Pol II.

**Figure 2.**
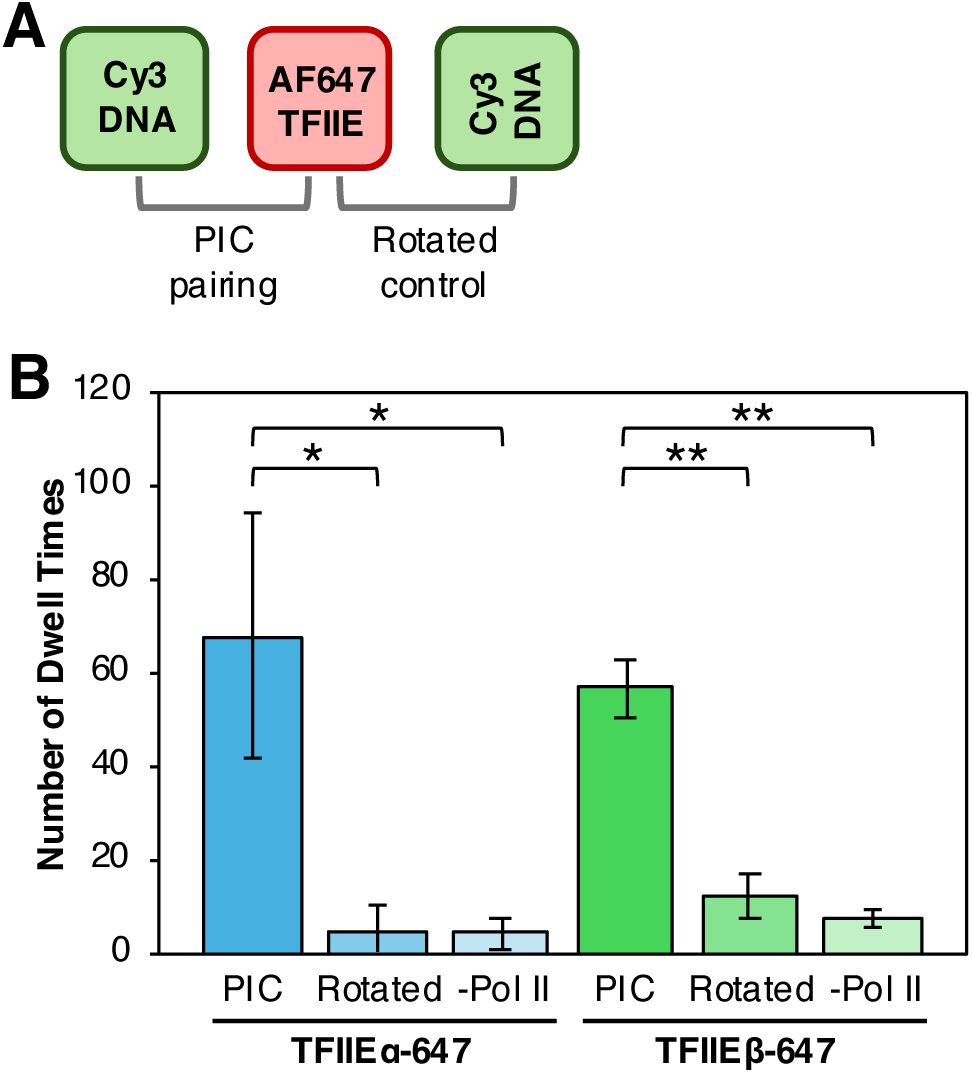
TFIIE-647 dynamics are dependent on Pol II and occur at colocalized spot pairs. A) In the rotated control analysis, the green Cy3 DNA movie was rotated 90° counterclockwise, then used for spot pairing and image analysis. B) Plotted are the average number of dwell times obtained after colocalization in PICs, rotated controls, and excluding Pol II from PIC assembly. Error bars are the standard deviation (n=4, PIC and rotated control; n=3,-Pol II). p-values are from a two-sided unpaired t test, *p ≤ 0.01, **p ≤ 0.001.

### TFIIEα-647 and TFIIEβ-647 dynamically associate with PICs

Using the system described above, we first investigated the interaction of TFIIEα with PICs in real time. Four experimental replicates were conducted, the unbound and bound dwell times were pooled for all replicates, then plotted as a cumulative sum. The plots were fit with exponential equations to obtain association and dissociation rate constants. Unbound dwell times fit best with a single exponential to give k_on(obs)_, indicating a single kinetic population during association (Figure 3A, left plot). This is an observed association rate constant because the measurements were made at a single concentration of labeled protein. Bound dwell times fit best with a double exponential to give k_off_1_ and k_off_2_, indicating two kinetic populations governing dissociation (Figure 3A, right plot). Rate constants are shown on their respective plots and in Table 1, which also includes the 95% confidence intervals of the curve fits, half-times (t_1/2_), and the population percentages associated with each k_off_ value.

**Figure 3.**
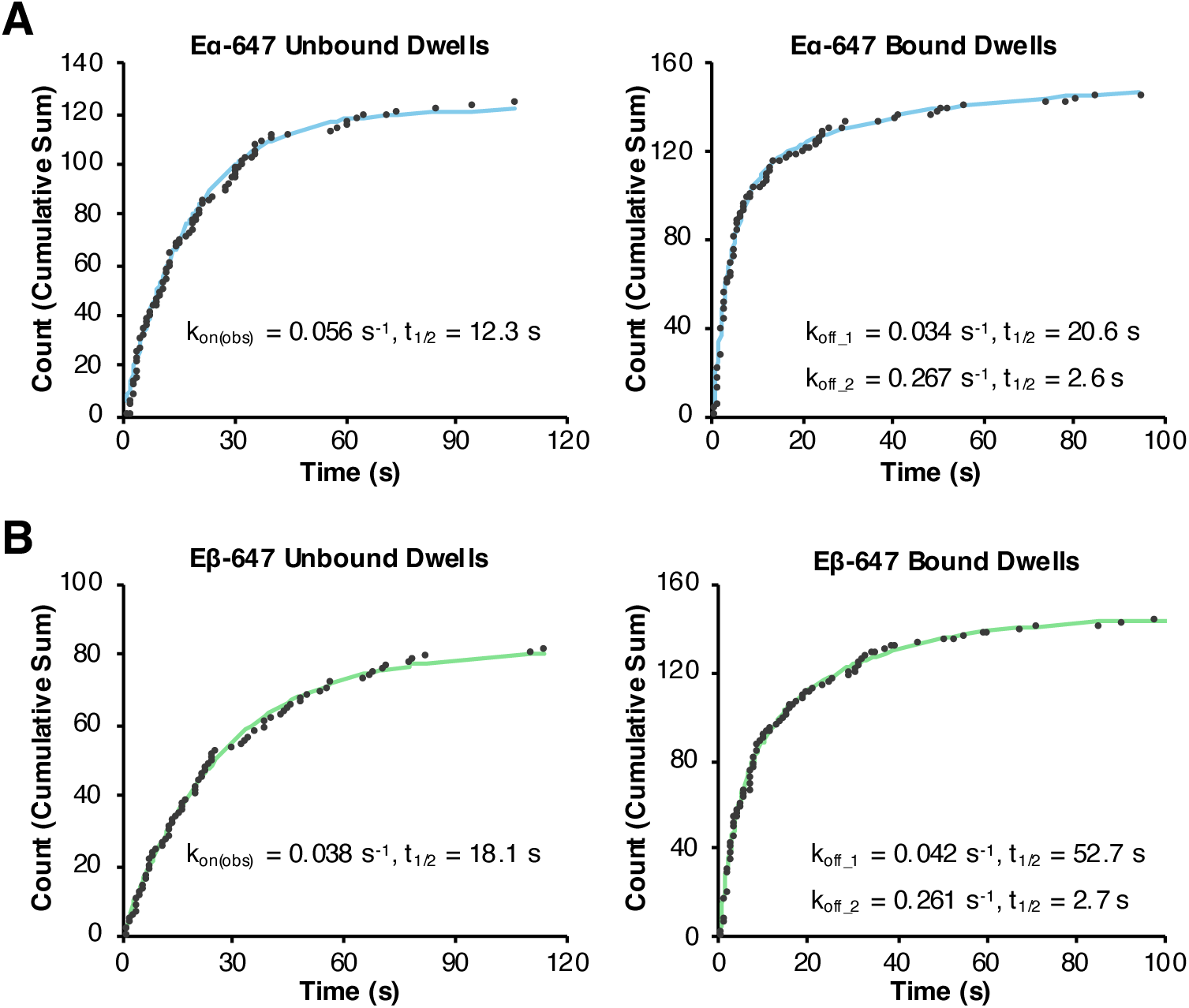
TFIIEα and TFIIEβ show dynamic interactions with PICs. A and B) Cumulative sum plots and rate constants for TFIIEα-647 and TFIIEβ-647, respectively. Unbound dwell times were plotted in cumulative sum plots and fit with single exponential equations, producing the association rate constants (k_on(obs)_) and half-times (t_1/2_) shown. Bound dwell times were plotted in cumulative sum plots and were best fit with double exponential equations, producing the dissociation rate constants (k_off_1_ and k_off_2_) and half-times (t_1/2_) shown. Dwell times were collected from four experimental replicates each for TFIIEα and TFIIEβ.

**Table 1.** Rate constants for TFIIEα-647 and TFIIEβ-647 association and dissociation from PICs in the presence and absence of TFIIH. Data from four independent replicates were combined and fit with exponential equations to obtain the rate constants shown. The 95% CI is the confidence interval of the rate constant obtained from the curve fit. Half-times were calculated using t_1/2_ = ln(2)/k. Pop % is the percentage of the population represented by each rate constant in a double-exponential fit.

| Protein | Rate Constant | Complete PIC |  |  |  | Excluding TFIIH |  |  |  |
| --- | --- | --- | --- | --- | --- | --- | --- | --- | --- |
|  |  | k (s<sup>-1</sup>) | 95% CI | t<sub>(1/2)</sub> | Pop % | k (s<sup>-1</sup>) | 95% CI | t<sub>(1/2)</sub> | Pop % |
| TFIIα | k<sub>on(obs)</sub> | 0.056 | 0.051 - 0.062 | 12 | - | 0.080 | 0.077 - 0.083 | 8.7 | - |
|  | k<sub>off_1</sub> | 0.034 | 0.026 - 0.043 | 21 | 32 | 0.116 | 0.108 - 0.123 | 6.0 | 33 |
|  | k<sub>off_2</sub> | 0.27 | 0.23 - 0.31 | 2.6 | 67 | 0.50 | 0.47 - 0.54 | 1.4 | 67 |
| TFIIβ | k<sub>on(obs)</sub> | 0.038 | 0.035 - 0.042 | 18 | - | 0.06 | 0.05 - 0.08 | 11 | - |
|  | k<sub>off_1</sub> | 0.042 | 0.038 - 0.046 | 17 | 52 | 0.030 | 0.028 - 0.032 | 23 | 55 |
|  | k<sub>off_2</sub> | 0.26 | 0.22 - 0.31 | 2.7 | 47 | 0.31 | 0.26 - 0.39 | 2.2 | 45 |

The same methods were applied to investigate TFIIEβ-647 dynamics during PIC assembly: four replicates were performed, then pooled dwell times were plotted as a cumulative sum and fit with exponential equations. The rate constants are displayed on the plots in Figure 3B and listed in Table 1 with 95% confidence intervals, population percentages, and half-times (t_1/2_). Like TFIIEα-647, TFIIEβ-647 also exhibited one kinetic population for association (k_on(obs)_ obtained from unbound dwells) and two kinetic populations for dissociation (k_off_1_ and k_off_2_ obtained from bound dwells). These data provide clear evidence that both subunits of TFIIE exhibit dynamic on/off binding with PICs.

To compare the kinetics of TFIIEα-647 and TFIIEβ-647, curve fits from their respective cumulative sum plots were normalized to 1.0 and plotted together (Figure 4A and B, left plots). These comparisons are also shown in bar plots of association and dissociation rate constants, with the error bars showing the 95% confidence intervals (Figure 4A and B, right plots). TFIIEα (blue) showed an association rate constant 1.5-fold greater and a half-time 1.5-fold shorter than TFIIEβ. This difference in k_on(obs)_ is reflected in the dissimilar curve fits for their respective data sets and the bar plot with non-overlapping 95% confidence intervals. By contrast, the k_off_1_ and k_off_2_ dissociation rate constants were similar for TFIIEα and TFIIEβ and had overlapping confidence intervals, suggesting similar kinetic stability in PICs. Interestingly, despite similar rate constants, the bound dwell curve fits are visually dissimilar (Figure 4B, left plot); this is due to the difference in population distribution attributed to each rate constant for the two subunits (see Table 1). Taken together, these data show that the two subunits exhibit unique association kinetics and similar dissociation kinetics, suggesting asynchronous assembly into the PIC is possible.

**Figure 4.**
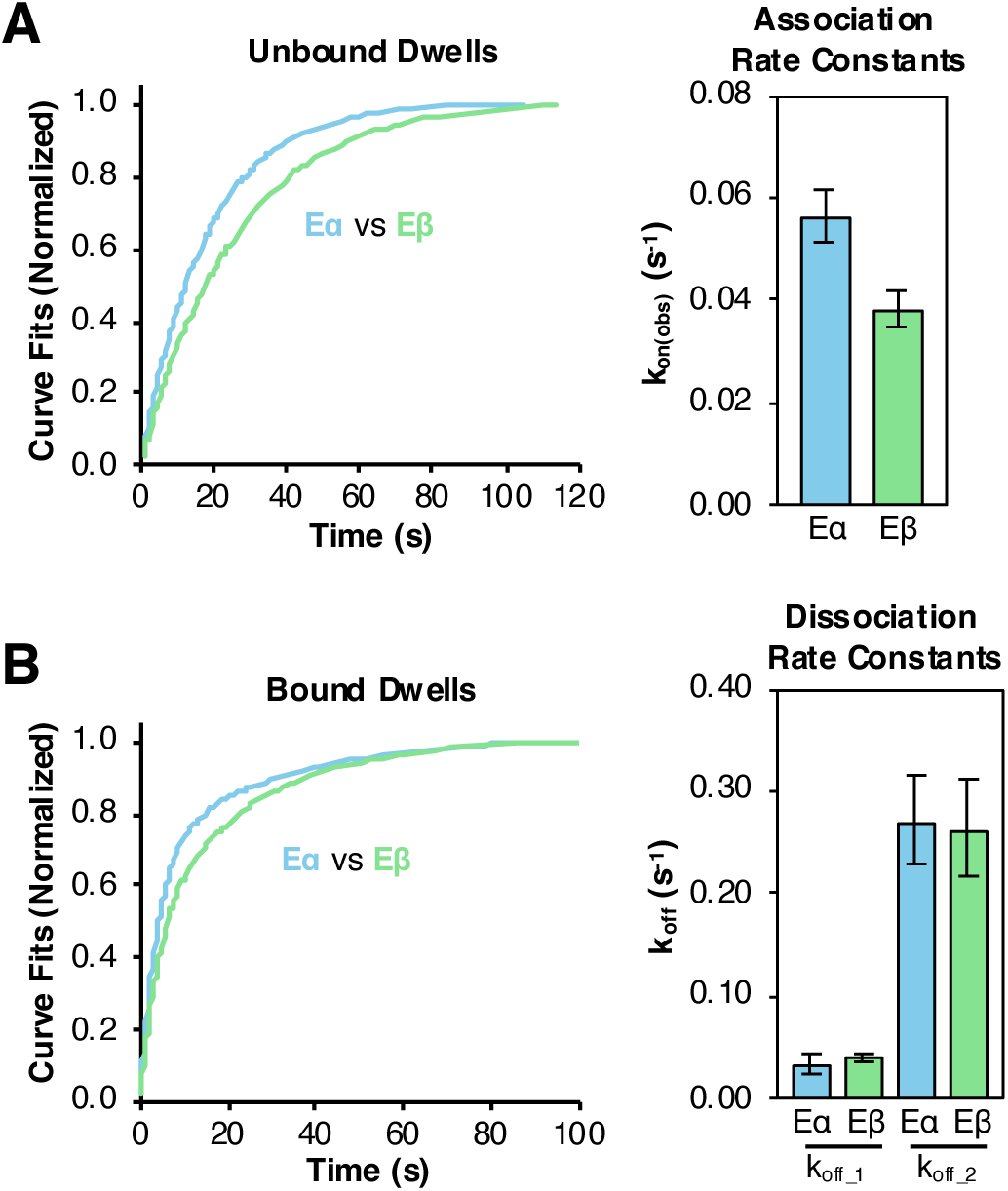
Comparing rate constants for TFIIEα and TFIIEβ association/dissociation with PICs shows unique kinetic behaviors for the two subunits. A and B) The curve fits for TFIIEα-647 (blue) and TFIIEβ-647 (green) were normalized to 1.0 for display on the same plot. The bar graphs show the rate constants obtained from the unbound (k_on(obs)_) and bound (k_off_1_, k_off_2_) dwells, with the error bars showing the 95% confidence intervals of the curve fits (n=4 for all data sets).

### TFIIH exclusion increases TFIIE dynamics, with a stronger effect on TFIIEα

We investigated how TFIIE dynamics changed when TFIIH was excluded from PICs. During PIC assembly, TFIIE recruits TFIIH to the PIC and stimulates the enzymatic activities of TFIIH [25–27,48,49]. TFIIH makes extensive contacts with other PIC proteins and the DNA downstream of the TSS, which could also serve to anchor TFIIE within the PIC [32,50]. Therefore, we hypothesized that omission of TFIIH would destabilize TFIIE in PICs because the additional network of protein-protein and protein-DNA interactions mediated by TFIIH would be absent. To test this, we measured rate constants for on/off binding of TFIIEα-647 and TFIIEβ-647 with PICs assembled in the absence of TFIIH (-H). Cumulative sum plots are shown in Figure 5A for TFIIEα-647 and in Figure 5B for TFIIEβ-647. The rate constants obtained from the curve fits are shown on the plots and in Table 1. Figures 5A and B also show bar plots of the average number of dwell times per experiment in the +TFIIH and-TFIIH conditions. Notably, we found that excluding TFIIH led to a ∼6-fold increase in the number of dwell times per experiment for TFIIEα-647 (Figure 5A, bar plot at right). This resulted from both a ∼3-fold increase in the number of on/off binding events per PIC and twice as many PICs with dynamic TFIIEα-647 molecules. By contrast, TFIIH exclusion did not change the average number of TFIIEβ-647 dwell times observed (Figure 5B, bar plot at right). Additionally, TFIIH exclusion did not impact the number of TFIIEβ-647 on/off binding events per PIC.

**Figure 5.**
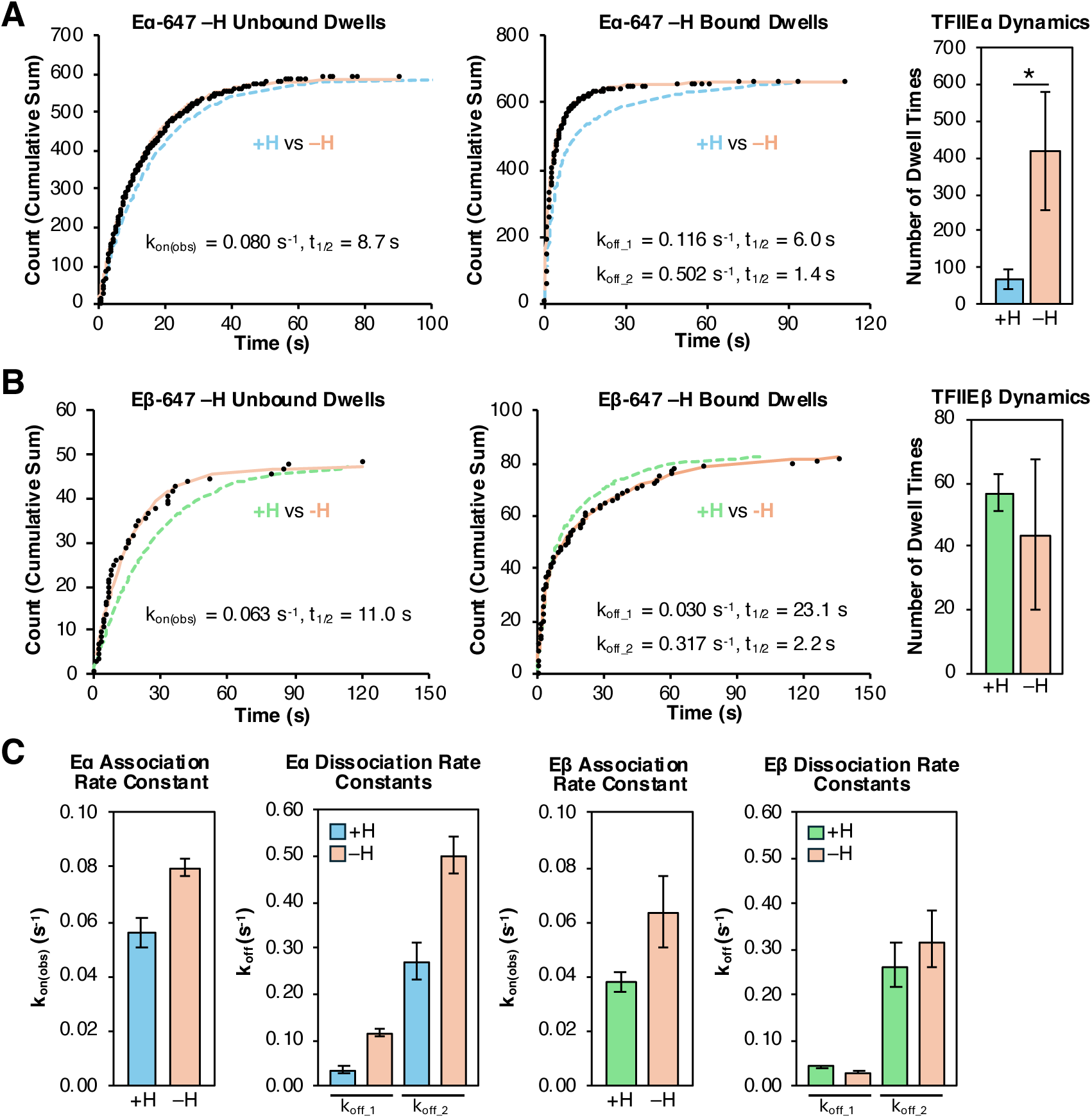
Exclusion of TFIIH from PIC assembly changes the dynamics of TFIIE, with a stronger effect on TFIIEα compared to TFIIEβ. A) Unbound and bound dwells for TFIIEα-647 were plotted on cumulative sum plots and fit with exponential equations (orange) to obtain the rate constants and half-times shown on the graphs. The blue dashed lines (+TFIIH) are the curve fits from panel 3A. The bar graph on the right compares the average number of dwell times when including (blue) and excluding (orange) TFIIH; the error bars are the standard deviation (n=4 for +TFIIH, n=3 for-TFIIH). p-value from a two-sided unpaired t test was *p = 0.007. B) Unbound and bound dwells for TFIIEβ-647 were plotted on cumulative sum plots and fit with exponential equations to obtain the rate constants and half-times shown on the graphs. The green dashed lines (+TFIIH) are the curve fits from panel 3B. The bar graph on the right compares the average number of dwell times when including (green) and excluding (orange) TFIIH; the error bars are the standard deviation (n=4, +TFIIH; n=3,-TFIIH). C) Bar graphs comparing rate constants when including (blue and green) and excluding (orange) TFIIH from PIC assembly for each TFIIE subunit; error bars are 95% confidence intervals.

Figure 5C shows comparisons of the rate constants across the +TFIIH (blue and green bars for TFIIEα and TFIIEβ, respectively) and-TFIIH (orange bars) conditions. In nearly every case (except for TFIIEβ k_off_1_), omitting TFIIH caused an increase in the rate constant. Additionally, all rate constant comparisons (except for TFIIEβ k_off_2_) do not have overlapping 95% confidence intervals, showing that excluding TFIIH from PICs strongly impacts the kinetics of on/off binding for the TFIIE subunits. For TFIIEα-647, all three rate constants increased in the-TFIIH condition, showing that both association and dissociation occur more rapidly. The strongest effect was on dissociation, reflecting a loss in kinetic stability. In the absence of TFIIH, k_off_1_ and k_off_2_ for TFIIEα-647 increased by 3.4-fold and 1.9-fold, respectively. Accordingly, the calculated half-times also significantly decreased, from 21 s to 6.0 s (k_off_1_) and 2.6 s to 1.4 s (k_off_2_).

For TFIIEβ-647, we also observed faster association kinetics in the absence of TFIIH. Indeed, k_on(obs)_ increased 1.7-fold and the association half-time decreased from 18 s to 11 s. By contrast, the dissociation rate constant k_off_1_ decreased by 1.4-fold in the absence of TFIIH, while k_off_2_ wasn’t significantly changed, indicating TFIIH omission slightly stabilized TFIIEβ. Together our data show that loss of TFIIH significantly destabilized TFIIEα in PICs, increasing both the rate and frequency of on/off binding events. The impact of TFIIH exclusion on TFIIEβ was smaller in magnitude. Overall, our real-time kinetic measurements support the model that TFIIE stability within PICs is strengthened by protein-protein and protein-DNA interactions facilitated by TFIIH.

### TFIIEβ point mutations associated with disease affect the kinetics of TFIIE assembly into PICs

In humans, mutations in the TFIIEβ subunit have been linked to the rare genetic disorder trichothiodystrophy (TTD). Mutations in various subunits of TFIIH can also give rise to TTD, with these patients experiencing increased sensitivity to UV light (xeroderma pigmentosum) and impaired nucleotide-excision repair [51,52]. TTD patients with TFIIEβ mutations and wild type TFIIH (XPD subunit) do not have photosensitivity or impaired nucleotide excision repair; this is typically designated as the non-photosensitive variant of TTD (NPS-TTD). Two different point mutations in TFIIEβ are responsible for NPS-TTD: A150P and D187Y [53,54]. Both mutations occur in the winged-helix 2 (WH2) domain of TFIIEβ, which is critical for the association of TFIIEβ with TFIIEα and the p62 subunit of TFIIH [55]. CryoEM analyses suggest that each of the mutations destabilize the secondary structure packing of the WH2 domain [55]. Additionally, these mutations impair transcription initiation in vitro, as shown by abortive initiation assays [45]. We tested the activity of AF647-labeled TFIIEβ mutants A150P and D187Y in our purified in vitro transcription system. As shown in Figure 6A, the fractional activity of both mutants (lanes 4 and 5) is lower than the wild type protein (lane 1), as expected from previous literature [45].

**Figure 6.**
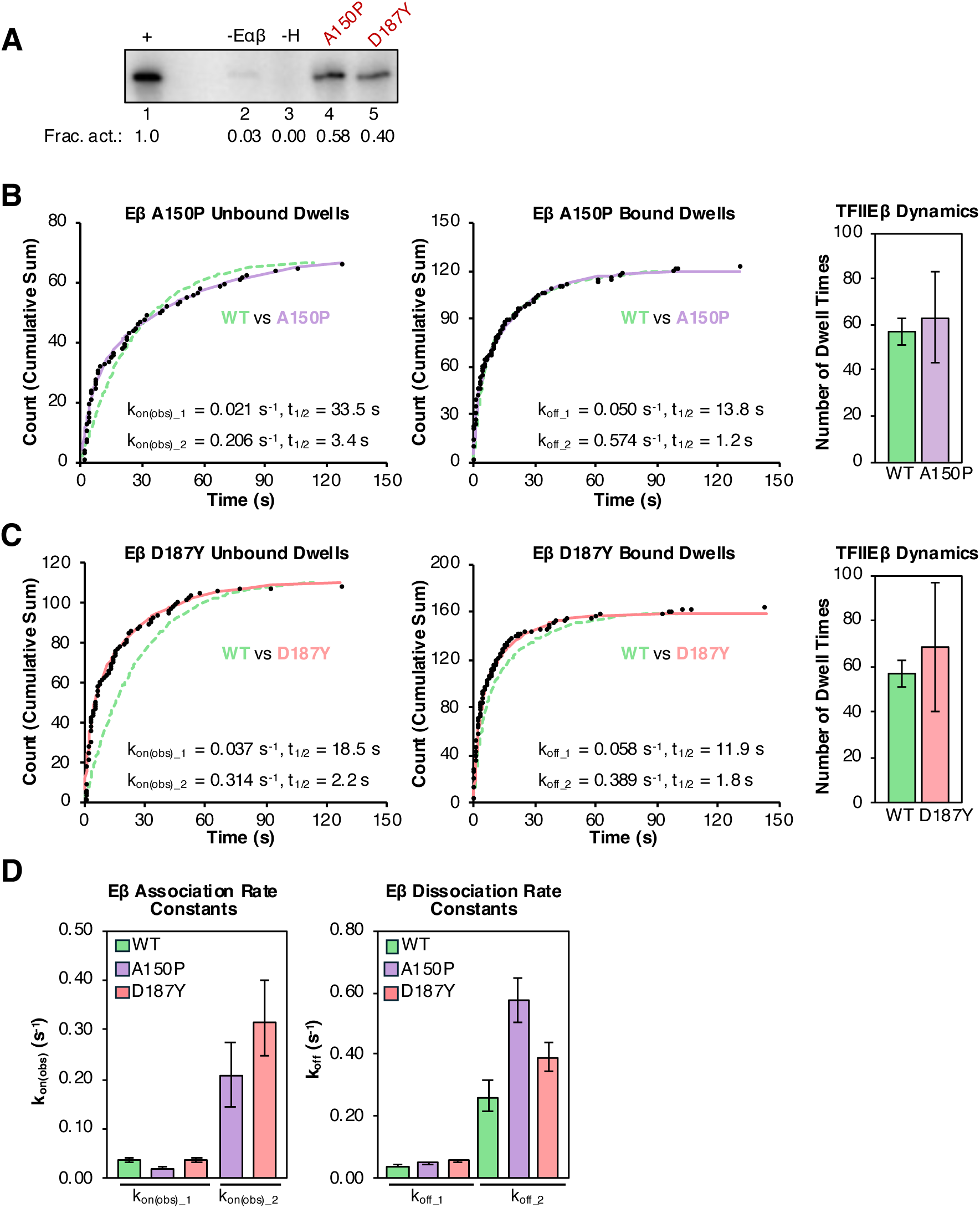
Point mutations in TFIIEβ impact its kinetic behaviors. A) AF647-labeled TFIIEβ mutants (A150P and D187Y) show lower levels of transcriptional activity than their WT counterparts. Shown is ^32^P-labeled RNA from ensemble in vitro transcription assays. Fractional activity was quantified compared to the positive control in lane 1. A well was skipped between lanes 1 and 2. B) Unbound and bound dwells for TFIIEβ-647 A150P were plotted on cumulative sum plots and fit with exponential equations, with rate constants and half-times shown on the graphs. The curve fits for WT TFIIEβ (green dashed line) were normalized to the curve fits for A150P (purple) and plotted on the same graphs for comparison. The bar graph on the right plots the average number of dwell times for wild type and A150P TFIIEβ, the error bars are the standard deviation (n=4 for WT, n=3 for A150P). The WT data was from the experiments shown in Figure 3B. C) Unbound and bound dwells for TFIIEβ-647 D187Y were plotted on cumulative sum plots and fit with exponential equations, with rate constants and half-times shown on the graphs. The curve fits for WT TFIIEβ (green dashed line) were normalized to the curve fits for D187Y (coral line) and plotted on the same graphs for comparison. The bar graph on the right plots the average number of dwell times for wild type and D187Y TFIIEβ, the error bars are the standard deviation (n=4 for both WT and D187Y). The WT data was from the experiments shown in Figure 3B. C) Bar graphs comparing rate constants for WT Eβ (green), Eβ A150P (purple), and Eβ D187Y (coral); error bars are 95% confidence intervals.

Using our smTIRF system, we explored the effects of the A150P and D187Y mutations on the on/off binding kinetics of TFIIEβ. With TFIIEβ-A150P, there was no significant difference in the frequency of dynamic binding events (Figure 6B, bar plot at right) compared to the wild type protein (WT); however, the rates of on/off binding changed, particularly for association events. The rate constants obtained from the curve fits of dwell times are shown on the plots in Figure 6B and in Table 2. Whereas the unbound dwells for WT TFIIEβ showed one kinetic population for association, unbound dwells for TFIIEβ-A150P were best fit with a double exponential, giving rise to two kinetic populations (Figure 6B). The k_on(obs)_1_ and k_on(obs)_2_ association rate constants for TFIIEβ-A150P were 1.8-fold lower and 5.4-fold greater (respectively) than the single k_on(obs)_ for WT TFIIEβ (Table 2). The new faster kinetic population that appeared accounted for 38% of the total population. For dissociation, TFIIEβ-A150P showed faster kinetics than WT TFIIEβ, with k_off_1_ and k_off_2_ rate constants 1.2-fold greater and 2.2-fold greater than the WT k_off_ values, respectively (Figure 6B, middle plot; the purple A150P line largely overlaps the green WT line for bound dwells). Overall, the decrease in kinetic stability (i.e. increased k_off_ values), and the appearance of a rapid second population for association suggest a destabilization of TFIIEβ within the PIC due to the A150P mutation. We investigated the effects of the D187Y mutation on TFIIEβ dynamics in the same manner. The frequency of TFIIEβ-D187Y on/off binding events was similar compared to WT TFIIE (Figure 6C, bar plot at right). Like TFIIEβ-A150P, TFIIEβ-D187Y also showed a new kinetic population for association. The k_on(obs)_1_ association rate constant for TFIIEβ-D187Y was nearly identical to the k_on(obs)_ for WT Eβ, but k_on(obs)_2_ was 8.2-fold greater (Figure 6C and Table 2). The new faster kinetic population that appeared accounted for 43% of the total population. The k_off_1_ and k_off_2_ dissociation rate constants for TFIIEβ-D187Y were slightly (1.4-fold and 1.5-fold) greater than those for WT TFIIEβ, respectively. The bar plots in Figure 6D compare the rate constants between the mutant and WT TFIIEβ proteins. When comparing the two mutants, it appears that the A150P mutation had a larger impact on dissociation kinetics, leading to greater destabilization. Both the A150P and D187Y mutations had a large impact on association, leading to new kinetic populations that bound PICs faster than the WT protein.

**Table 2.**
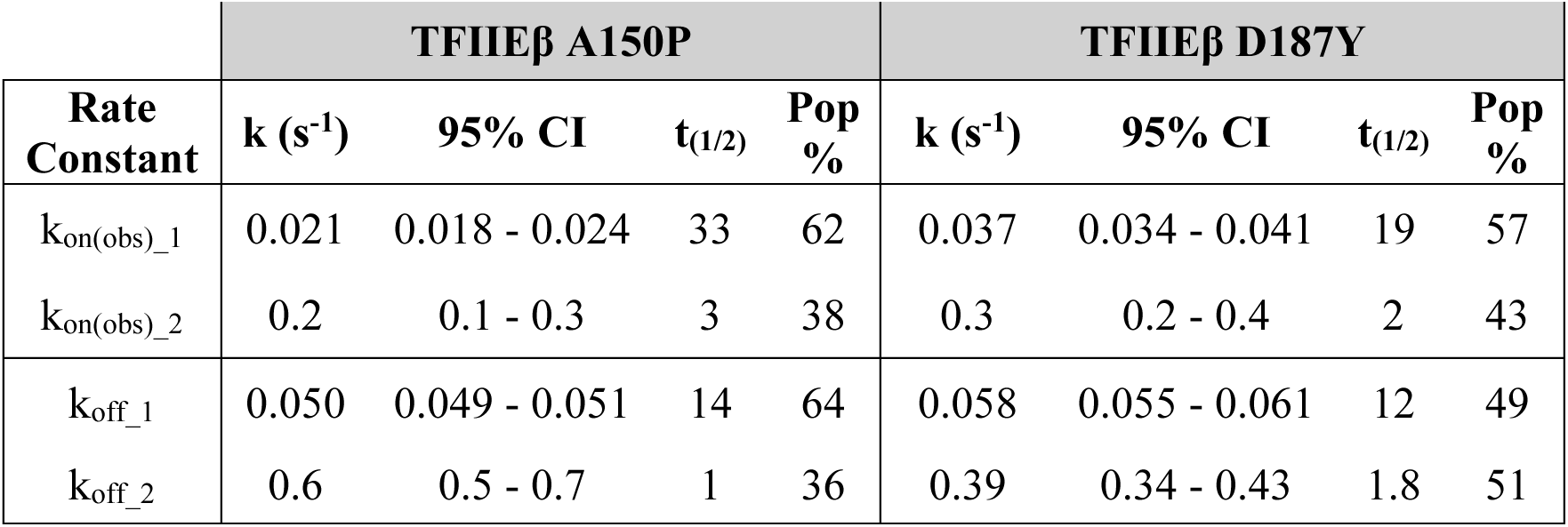
Rate constants for two mutant TFIIEβ-647 proteins (A150P and D187Y) associating and dissociating from PICs. Data from four independent replicates were combined and fit with exponential equations to obtain the rate constants shown. The 95% CI is the confidence interval of the rate constant obtained from the curve fit. Half-times were calculated using t_1/2_ = ln(2)/k. Pop % is the percentage of the population represented by each rate constant in a double-exponential fit.

## Discussion

The Pol II-specific GTFs facilitate important functions during early transcription: PIC assembly, opening of the promoter DNA, phosphorylation of the Pol II CTD, initiation, and early transcription. The coordination of the GTFs provides multiple layers of transcriptional regulation, and understanding these regulatory mechanisms is necessary for understanding general mechanisms of Pol II transcription. General processes governing PIC assembly, including the order in which GTFs bind to the PIC (in the stepwise model of assembly) have been described [2,4,5], but information about the kinetics and dynamics governing individual factors within PICs is lacking. With the goal of gaining this insight, our lab has developed a fluorescence-based smTIRF system to investigate the dynamics of individual GTFs in PICs. Here we investigated the dynamics of TFIIEα and TFIIEβ, which serve to recruit and regulate the enzymatic activities of TFIIH. We observed highly dynamic behavior from both TFIIE subunits during PIC assembly, which was dependent on Pol II. We also found that excluding TFIIH altered the dynamics of both TFIIE subunits, significantly destabilizing TFIIEα. Lastly, we also found that two disease-associated TFIIEβ point mutations showed more rapid kinetic behaviors in PICs compared to WT TFIIEβ, perhaps explaining their negative impact on transcriptional activity.

Our real-time imaging showed that both TFIIE subunits exhibited dynamic interactions with the PIC, with multiple binding and unbinding events seen throughout our imaging time course of 150 seconds. These results support the emerging model that PICs are not static assemblies. Dynamic association of TFIIE with PICs in yeast extracts has also been observed using in vitro single-molecule imaging techniques [23]. Repeated cycles of yeast TFIIEα (Tfa1) association and dissociation were observed during a single TFIIF-Pol II association event occurring during PIC assembly. Interestingly, the dynamic behavior of the TFIIE subunits directly contrasts with the documented behavior of other GTFs such as TFIIB and TFIIF, which show stable binding to the PIC or few dynamic exchanges [23,24]. This suggests that TFIIE dynamics are unique and could play a regulatory role in setting the activity of PICs as they move through the transcription cycle.

TFIIEα and TFIIEβ showed similar kinetics for dissociation, but unique kinetics for association, suggesting that asynchronous assembly may be taking place. This is interesting, given that TFIIE is canonically thought to bind to the PIC as a heterodimer of αβ subunits [25–28]. Previous literature suggests an asynchronous model of release [45]. An ensemble transcription study found that TFIIEα and the CAK domain of TFIIH released from PICs after Pol II CTD phosphorylation, while TFIIEβ and the core domain of TFIIH released after initiation, as elongation factors were recruited [45].

The exclusion of TFIIH produced a striking increase in dynamic behavior, in particular for TFIIEα. Both subunits showed an increase in their association rate constants and most of the dissociation rate constants, indicating faster binding and release from PICs when TFIIH is absent. In addition, a large effect on the frequency of TFIIEα association and dissociation was observed; the average number of dynamic events increased 6-fold in the absence of TFIIH. This was not observed with TFIIEβ. This suggests that the absence of TFIIH increases the frequency with which TFIIEα monomers (i.e. TFIIEα not associated with TFIIEβ) bind to and release from PICs. In other words, TFIIH prevents TFIIEα monomers from binding PICs. Together, the changes in the rate constants and the number of dynamic on/off events suggest that TFIIH stabilizes the binding of the TFIIE heterodimer to the PIC. This is likely due to the extensive network of intermolecular contacts facilitated by TFIIH with both downstream DNA and the other factors in the PIC. This idea is supported by structural studies using cryo-EM, which noted difficulties resolving TFIIE in the absence of TFIIH unless TFIIE was added in excess, suggesting TFIIE interaction with PICs in the absence of TFIIH was highly dynamic [35,38]. These results reveal kinetic behaviors that refine our understanding of PIC assembly.

For each subunit of TFIIE we observed one kinetic population for association (k_on(obs)_) and two kinetic populations for dissociation (k_off_1_ and k_off_2_). These observations are consistent with the model that some PICs experience a conformational change after TFIIE binding, which increases the stability of TFIIE within the PIC. The faster kinetic population we observed for dissociation may approximate the rate of release of TFIIE from PICs before the stabilizing conformational change. In stabilized complexes, release of TFIIE would be slower, perhaps undergoing two steps for dissociation. The difference between the fast and slow dissociation rate constants we measured range from 5-fold to 10-fold; this large difference suggests that TFIIE stability is highly dependent on this conformational shift. This model is consistent with the data for each subunit and may also approximate the behavior of the αβ heterodimer; determining this will require further experimentation.

We also investigated the impacts of two clinically significant point mutations in TFIIEβ: A150P and D187Y. While wild type TFIIEβ showed a single observed association rate constant, both mutants showed two distinct kinetic populations binding to PICs. Molecules in the new kinetic population (∼40% of the total) associated faster, with an observed rate constant over 4-fold higher than the single k_on(obs)_ seen for wild type TFIIEβ. It is possible that the rapid populations arise from new, transient interactions with PIC components that only occur with the mutant proteins. We also saw an increase in both dissociation rate constants for both mutants, indicating decreased stability of the mutants within the PIC. This change in stability may explain the lower fractional activity that we observed in our ensemble activity assays compared to wild type TFIIEβ.

The advent of single-molecule imaging has enabled exploration of the dynamic interactions governing the many moving parts of the Pol II transcription system, including the GTFs and Pol II. Here we have revealed that TFIIE is a dynamic component of the PIC, its dynamics can be regulated by the binding of TFIIH, and TFIIEβ mutations which destabilize TFIIEβ within the PIC may impact activity. Ultimately, it is our goal to connect the dynamics of TFIIE to the fates of complexes as they transcribe.

## Methods

### Design and synthesis of transcription DNA templates

The sequence of the DNA template is as follows: 5’CTCTAGAGGATCCCCGGTGTTCCTGAAGGGGGGCTATAAAAGGGGGTGGGGGCG CGTTCGTCCTC**ATCACTATCTTTAATCACTACTCACACTAACCTCACC**ACCCTACTCTCCCTT CCCTATCCCTTATCTTAACCACTCCAATTACATACACCTTCTTCTATATTTCCCAAATCTATCATCATTCACTCTCATCCCCTCTTCCTTCACTCCCATTCTATTCTACTCCTTTCCCTTTCCATATCCCCTCCACCCCCCTTCCTCCCCTCTTTCAATCTTATCCCCAATCATAAAATTATCTCAATTATATTCTCCTTCCATACCCCCTATCATCCTCATCCCTATCACCCCCTACTCACCCAATACTCCTCACACTC ATTTCTCATTCCACTCCC**GGG**-3’

The DNA template contained the AdMLP (-40 to-1, <u>underlined</u>) and the early transcribed sequence was previously optimized for template usage (+1 to +37, **bold**) [47]. The transcribed region contained a G-less cassette from +1 to +330, followed by a GGG roadblock (3’ end, **bold**). The sequence was inserted into a pBS-KS+ plasmid backbone using HindIII and EcoRI. DNA templates used in TIRF experiments were synthesized using PCR; the template strand reverse primer contained a 5’-biotin and a Cy3 on the adjacent base (IDT). Amplified DNA was purified using a Monarch PCR & DNA Cleanup Kit (NEB).

### Protein expression and purification

Genes for TFIIE constructs were cloned into a pET-21a+ vector using NdeI and BamHI. Cloning yielded two final gene products: 2xHA-TFIIEα-GSSGGSSG-SNAP-6xHis and TFIIEβ-6xHis. The two TFIIEβ mutants (A150P and D187Y) were created using site-directed mutagenesis and the aforementioned TFIIEβ-6xHis construct.

After expression in BL21 E. coli, cells were pelleted then lysed by sonication in lysis buffer (20 mM Tris pH 7.9, 5 mM BME, 250 mM NaCl, 10% glycerol, 1X Protease Inhibitor cocktail (Roche), 200 µM PMSF, and 10 mM imidazole), then centrifuged again to pellet cell debris. The lysis supernatant was then loaded onto a gravity column containing 1.3 mL of HisPur Ni-NTA resin (Thermo-Fisher), which was equilibrated with 7 mL of lysis buffer. After loading the supernatant onto the column, the column was washed with 7 mL of lysis buffer followed by 9 mL of wash buffer (20 mM Tris pH 7.9, 5 mM BME, 250 mM NaCl, 10% glycerol, 1X Protease Inhibitor cocktail (Roche), 200 µM PMSF, and 50 mM imidazole). Protein was eluted from the column with 5 mL of elution buffer (20 mM Tris pH 7.9, 5 mM BME, 250 mM NaCl, 10% glycerol, 1X Protease Inhibitor cocktail (Roche), 200 µM PMSF, and 250 mM imidazole), and collected in 1 mL fractions. SDS-PAGE was used to determine the location of protein in the elution fractions. Chosen eluate fractions were pooled and dialyzed overnight at 4°C in dialysis buffer (10 mM Tris pH 7.9, 1 mM DTT, 50 mM KCl, and 10% glycerol), then aliquoted in batches of 200 µL for fluorescent labeling. For TFIIEβ labeled using maleimide chemistry, DTT was excluded from the dialysis buffer.

In addition to the C-terminal 6xHis tag, TFIIEα also contained two copies of an N-terminal HA antibody affinity tag (sequence of MGYPYDVPDYAIEGRH per tag) for further purification. Anti-HA antibody (suspended in PBS and 20 mM Tris pH 7.9) was conjugated to protein A Sepharose beads for purification. 70 µL of protein A Sepharose Fast Flow beads (GE Healthcare) was added to a tube on ice. The beads were washed four times with 10 bead volumes (700 µL) of wash buffer (1X PBS + 0.1% NP-40). With each wash, beads were centrifuged at 800 x g for 2 mins, then the wash buffer was removed. After the final wash, 500 µL of 12CA5 antibody solution was added, then nutated overnight at 4°C to conjugate the antibody to the beads. After conjugation, the beads were centrifuged for 2 mins, then washed once with 10 bead volumes of TGEMD(0.2) (20 mM Tris pH 7.9, 20% glycerol, 1 mM EDTA, 5 mM MgCl_2_, 200 mM NaCl, 0.1% NP-40, 200 µM PMSF, 1X Protease Inhibitor cocktail (Roche), and 1 mM DTT), then once with TGEMD(1.0) (20 mM Tris pH 7.9, 20% glycerol, 1 mM EDTA, 5 mM MgCl_2_, 1 M NaCl, 0.1% NP-40, 200 µM PMSF, 1X Protease Inhibitor cocktail (Roche), and 1 mM DTT).

TFIIEα was first purified using the Ni-NTA protocol, then the eluate (after dialysis) was combined with the antibody-conjugated beads and nutated at 4°C for 2.5 hours. The beads were then washed twice with 10 bead volumes of TGEMD(1.0), then twice with 10 bead volumes of TGEMD(0.2). To elute the protein from the beads, 200 µL of elution buffer (20 mM Tris pH 7.9, 20% glycerol, 1 mM EDTA, 5 mM MgCl_2_, 200 mM NaCl, 0.1% NP-40, 200 µM PMSF, 1X Protease Inhibitor cocktail (Roche), 1 mM DTT, and 1 mg/mL HA peptide) was added, then nutated with the beads at 4°C for 1.5 hours before centrifuging (800 x g for 2 min). After collecting the supernatant with the eluted protein, a second batch of 200 µL elution buffer was added, then incubated with the beads overnight before another elution. The two eluates were filtered separately through a Millex-GV4 filter to remove remaining beads, then snap-frozen in liquid nitrogen and stored at-80°C.

TBP was purchased from Active Motif (cat# 81114). TFIIB and TFIIF (as RAP30 and RAP74) were purified as previously described [56–58]. Pol II was isolated from HeLa cells as previously described [59]. TFIIH was purified as previously described [60].

### Fluorescent labeling of proteins

SNAP-tag labeling was used for purified TFIIEα: 100 µL of protein eluate and 4 µL of SNAP-Surface Alexa Fluor 647 substrate (10 µM final concentration resuspended in DMSO, NEB) was nutated at room temperature for 2 hours. Maleimide labeling was used for TFIIEβ: 200 µL of protein eluate was mixed with a 10X molar excess of TCEP and a 2X molar excess of maleimide AF647 dye substrate (resuspended in DMSO, VWR), then nutated at 4°C for 2.5 hours. After labeling reactions, excess free dye was removed using Pierce dye removal resin (Thermo Scientific). 40 µL of resin was used per 100 µL of labeling reaction. The resin was washed three times with 10 volumes of DB(150) buffer (10 mM Tris pH 7.9, 150 mM KCl, 10% glycerol, and 1 mM DTT). After the third wash, the protein dye labeling reaction was added to the resin, thoroughly mixed by pipetting, then immediately centrifuged (1,000 x g for 1 min at room temperature). The supernatant containing the dye-labeled protein was isolated, aliquoted (2-5 µL), snap-frozen in liquid nitrogen, then stored at-80°C.

### Testing labeled TFIIEα and TFIIEβ for activity using ensemble transcription assays

Ensemble transcription assays used the same purified DNA template from smTIRF experiments. Pol II and the GTFs were assembled in ensemble transcription reactions, as described before [46,47]. Each reaction (20 µL total volume) contained 2 nM of DNA. Minimal PICs were assembled (DNA, TBP, TFIIB, TFIIF, and Pol II) and incubated at room temperature for 10 mins, followed by addition of TFIIEα, TFIIEβ, and TFIIH and another incubation for 10 mins. After this, NTPs (625 µM ATP, 625 µM UTP, 91 µM CTP, 5 µCi [^32^P-α-CTP]) were added and the reactions incubated at room temperature for 30 mins. Reactions were quenched with 100 µL stop mix (3.125 M NH_4_OAc, 124.75 µg/mL yeast RNA, 150.0 µg/mL proteinase K, and 20 mM EDTA, syringe filtered with a 0.22 µm filter right before use) and ethanol precipitated to isolate transcribed mRNA. The radiolabeled mRNA transcripts were resolved using a denaturing gel and imaged using phosphorimagery.

### Single molecule TIRF experiments

Our objective-based single-molecule total internal reflection fluorescence (smTIRF) microscope was used for single-molecule experiments. Our system included a Nikon TE2000-U microscope, 1.49 NA immersion objective, and a Piezo nanopositioning stage. In the green channel, a 532 nm laser was used for excitation, and emission filters allowed 545-600 nm for imaging. In the red channel, a 640 nm laser was used for excitation, and emission filters allowed >660 nm for imaging. Images were collected using two Andor iXon Life EMCCD (IXON-L-897) cameras and NIS Elements software (Nikon Instruments).

Microscope slides and coverslips were thoroughly cleaned and functionalized with aminosilane, which allowed for conjugation of a mixture of mPEG and biotin-PEG to the surface. This process follows our previously published protocol [61]. For each experiment, a 150 µL mixture of 100 mM Trolox (Millipore Sigma) and 112 mM NaOH was made, vortexed, nutated for 1 hour at 4°C, then syringe filtered with a 0.22 µm filter. A cleaned, functionalized slide was washed twice with 200 µL of MiliQ water, then washed twice with 200 µL of DB/RM buffer (10 mM Tris pH 7.9, 50 mM KCl, 4 mM MgCl_2_, 10 mM HEPES pH 7.9, 10% glycerol, 1 mM DTT, and 0.05 mg/mL BSA). A streptavidin + BSA mix (0.2 mg/mL streptavidin and 0.8 mg/mL BSA, made with DB/RM in 50 µL total volume) was added and incubated for 5 mins, then the slide was washed twice with 200 µL of DB/RM. Four slide regions were then photobleached (∼1.5 mins per region) in the green channel.

PICs were assembled using one of two orders of addition: 1) DNA,TBP, TFIIB, TFIIF, Pol II; then TFIIE subunits and TFIIH, or 2) DNA; then TBP, TFIIB, TFIIF, Pol II; then TFIIE subunits and TFIIH. Using the first assembly method, RM buffer (20 mM HEPES pH 7.9, 8 mM MgCl_2_, and 1 mM DTT) was used to make a DNA mix consisting of 3-6 pM Cy3 DNA and 1 ng/uL dGdC competitor DNA (DNA: Poly(dG:dC) naked, InvivoGen). DB buffer (20 mM Tris pH 7.9, 100 mM KCl, 20% glycerol, 1 mM DTT, and 0.1 mg/mL BSA) was used to make the PIC protein mix consisting of 19.2 nM TBP, 5 nM TFIIB, 12 nM each of RAP30 and RAP74 (subunits of TFIIF), and 2-3 nM Pol IIO. 100 µL of a 50:50 mix of protein components in DB buffer and DNA components in RM buffer was made, gently mixed, incubated at room temperature for 5 mins, then added to the slide. Using the second assembly method, 3-6 pM of Cy3 DNA was added to a 50:50 mixture of RM and DB buffer, with a total volume of 100 µL. The DNA mixture was incubated on the slide for 5 mins; meanwhile, the minimal PIC factors (TBP, TFIIB, RAP30, RAP74, and Pol II) and dGdC competitor DNA were combined as described above, then incubated in a tube for 5 minutes at room temperature. The minimal PIC factors were added to the slide, then incubated for 10 mins. The EH mix consisted of 2-4 nM TFIIEα and TFIIEβ with one of the two subunits labeled with AF647, and 0.63 µL TFIIH (amount determined empirically from ensemble transcription assays) prepared in Imaging buffer (1 mg/mL glucose oxidase, 0.08 mg/mL catalase, 0.8% glucose, 3.45 mM Trolox made using DB/RM). Imaging Buffer (+EH) was added to the slide (100 µL) and incubated for 5 min before imaging.

The same four regions of each slide were imaged in the green and red channels. The green channel was first imaged for 50 frames at a 200 ms frame rate (10 seconds total) with a neutral density filter applied. The red channel was imaged for 750 frames at a 200 ms frame rate (150 seconds total), with red laser power of 124 mW, and with no neutral density filter applied. Images were analyzed using our in-house analysis software [61]. Frames for each movie were summed into a composite image, then green and red spots were colocalized across the two movies. While viewing the intensity traces for each channel, green-red spot pairs were manually sorted by the behavior of the red (TFIIE-647) dye into “stable” and “dynamic” categories. Dwell times were compiled for dynamic spots using our software, then plotted in cumulative sums plots in Microsoft Excel and fit with exponential curves to determine rate constants.

## Acknowledgements

This work was supported by grants MCB-2242824 from the National Science Foundation. S.A. was supported in part by training grant T32 GM008759 from the National Institutes of Health. We thank Anette Erbse and the Shared Instrument Pool core facility (RRID: SCR_018986) in the Department of Biochemistry at the University of Colorado Boulder for training and access to shared instrumentation. The Amersham ImageQuant 800 CCD (NIH, R35GM136392-04S1) and the Typhoon 5 (NIH, S10OD034218-01) were used for this project.

